# Characterization of bronchiolitis and vaccine-induced enhanced respiratory disease in Syrian hamsters caused by respiratory syncytial virus infection

**DOI:** 10.1101/2024.07.30.605875

**Authors:** Binbin Zhao, Gengxin Zhang, Jing Wu, Xiaoyue Ding, Xiaohui Wei, Wanjun Peng, Na Rong, Hekai Yang, Jiangning Liu

## Abstract

Respiratory syncytial virus (RSV) presents a significant health risk to pediatric and old populations. The quest for developing effective RSV vaccines and immunoprophylactic measures remains highly active, while the catastrophic enhanced respiratory disease (ERD) reaction triggered by RSV vaccines poses lingering concerns. Meanwhile, as the basis of vaccine research, the development of animal models of RSV infection has shown less substantial progress, with considerable disparities in immunological and pathological responses compared to humans. The Syrian hamster, which is reported to be susceptible to RSV infection, seems promising, but the lack of in-depth studies has hampered its application. Consequently, we have refined the RSV-infected hamster model by establishing infection models at different ages, reanalyzing virological and pathological data, and applying vaccination with heat-inactivated RSV virus for ERD reaction. In general, neonatal hamsters exhibited the highest level of viral amplification, while old hamsters displayed the most severe pathological manifestations, with adult hamsters in between. Although Syrian hamsters were moderately susceptible to RSV infection, they showed typical pathological changes of bronchitis in all age groups, with adult and old animals even showing clinical signs of coughing. In addition, hamsters demonstrated the typical ERD response to heat-inactivated virus, characterized by the exacerbation of lung injury with extensive neutrophil and eosinophil infiltration, as well as TH2 polarization. Collectively, these characteristics suggest that the Syrian hamster model has significant potential for investigating the pathogenesis of RSV infection, elucidating the mechanism of ERD, and evaluating vaccine efficacy and safety.

**Importance:** The Syrian hamster has been shown in our study to be a potential model for respiratory syncytial virus (RSV) infection, according to the following characteristics:

Hamsters of three different age groups all developed typical pulmonary bronchiolitis following RSV infection, whereas adult and old hamsters showed typical cough symptoms, comparable to those observed in clinical patients.

Hamsters of different ages showed considerable variability. Neonatal hamsters tolerated more viral replication, while old hamsters suffered the most severe pulmonary pathology.

Hamsters exhibited typical ERD responses after inoculation with heat-inactivated RSV virus, including enhanced lung pathology, pulmonary eosinophilic and neutrophil infiltration, and Th2 polarization, which was remarkable and more similar to the typical characteristics of the ERD response in humans.

## Introduction

The Respiratory syncytial virus (RSV), which is an enveloped, single-stranded, negative-sense RNA virus belonging to family *Pneumoviridae*, genus *Orthopneumovirus*, stands as a leading cause of respiratory illnesses, affecting individuals across the entire age spectrum [1]. While infections typically manifest as upper respiratory tract conditions, RSV poses a significant threat to vulnerable groups, such as children and the old, by potentially causing bronchiolitis and severe complications such as pneumonia, respiratory failure, and even death. Remarkably, RSV infects up to 90% of children by the time they reach two years of age and continues to reinfect older children and adults throughout their lives. According to 2022 Lancet data, RSV-related acute lower respiratory infections result in around 3.6 million hospitalizations and 26,300 deaths yearly among children aged 0-60 months [2]. In the United States, RSV is responsible for approximately 60,000 to 160,000 hospitalizations and 6,000 to 10,000 deaths annually among individuals aged 65 and above [3].

Since the discovery of RSV in the 1950s and its identification as a human pathogen, research has focused on the development of prophylactic vaccines and targeted therapeutic regimens. Despite the approval of two vaccines and a therapeutic antibody by the FDA, these still do not meet the preventive and therapeutic needs of all populations, particularly those at high risk [4–6]. Meanwhile, vaccine-induced enhanced respiratory syncytial virus disease (ERD), first observed in the 1960s following administration of formalin-inactivated RSV (FI-RSV) vaccine to children with no prior RSV exposure, is another challenge that must be considered in vaccine development [7].

Animal models of RSV infection are invaluable tools for the development of vaccines and therapies, as well as for the investigation of the immune mechanisms of infection. Currently, non-human primates (NHPs), rodents and ruminants are the most commonly used animal models. Because of the high specificity of human RSV to the human host, no other animal species can be naturally infected with human RSV, except for chimpanzees [8, 9]. However, the use of chimpanzees is extremely limited due to ethical and cost considerations. Other NHPs, including African green monkeys and Macaques, which are primarily employed to access vaccine efficacy and safety, exhibit semi-permissive replication of RSV, while the typical clinical features are absent [10–14]. Lamb models are effective in replicating the classic characteristics of RSV infection in preterm and neonatal infants [15, 16]. However, the maintenance and manipulation of lambs in experimental settings is challenging, and the availability of relevant research reagents is limited. The most widely used models remain those of rodents, including mouse and cotton rat models, both of which show pathological features such as bronchiolitis and mononuclear cell infiltration after RSV infection. Cotton rats have the advantage of being more sensitive to RSV and expressing antiviral set of Mx genes, while mice, as the most commonly used experimental animal, have obvious advantages, particularly with mature gene editing technology and various reagents to study the immune response. However, they also exhibit notable disparities in clinical symptoms, pathological alterations and immune responses to RSV infection when compared to humans [9, 17–23]. As there is no animal model that comprehensively reflects all aspects of this viral infection and disease, the strategy of developing different animal models to simulate as many aspects of RSV as possible is critical.

Due to its greater susceptibility to a much wider range of human pathogens, the Syrian hamster has been successfully used to establish a variety of animal models of infectious diseases, including SARS-CoV-2, Nipah virus, West Nile virus, Ebola virus, Yellow Fever virus and Influenza virus [24]. Despite the initial attempt to utilize the hamster as a model for RSV infection in 1970, subsequent studies limited in vaccine efficacy and antibody level assessment, neglecting an in-depth exploration of pathological changes and disease manifestation [25].

In this study, we performed infection experiments with RSV in three age groups of Syrian hamsters, recorded clinical symptoms, examined and analyzed viral replication and pathological changes to determine the optimal model parameters. We also indicated that the Syrian hamster model was capable of reproducing the ERD response by immunizing with inactivated RSV virus.

## Materials and methods

### Viruses and cell lines

Plaque-purified human RSV strain (RSV-A, long strain, ATCC) was propagated in Hep-2 cell lines utilizing Dulbecco’s Modified Eagle Medium (DMEM) enriched with 2% fetal bovine serum (FBS), penicillin (100 U/ml), and streptomycin (100 μg/ml). For the quantification of the 50% tissue culture infective dose (TCID_50_), viral suspensions were serially diluted in ten-fold increments and subsequently inoculated into Hep-2 cell cultures. Following a four-day incubation period, the determination of the endpoint dilution was based on the observation of cytopathic effects (CPEs) in 50% of the replicate cultures. The Reed-Muench method was employed to ascertain the viral titration, leveraging the endpoint dilution data.

### Animal experiment

The specific pathogen-free (SPF) neonatal (3 days old) and adult (6-8 weeks old) hamsters were purchased from Beijing Vital River Laboratory Animal Technology Co., Ltd. The SPF old hamster (70-80 weeks old) was obtained from Beijing HFK 263 Bioscience Ltd.

Hamsters were anaesthetized before intranasal infection for RSV infection. For dosing, each neonatal hamster received 10 μL of viral stock (10^7^ TCID_50_/ ml), while each adult and old hamster received 100 μl viral stock. For each age cohort, control groups were similarly administered an equivalent volume of phosphate-buffered saline (PBS) intranasally. Following inoculation, daily assessments were conducted to monitor weight fluctuations and clinical manifestations in the hamsters. Euthanasia was carried out on days 3, 5, and 7 to facilitate the collection of blood, lung, trachea, and nasal tissue samples, which were earmarked for further virological and histopathological analysis.

To elicit ERD responses in hamsters, RSV was inactivated by heating at 56 °C for 30 minutes. Subsequently, the inactivated virus was mixed with Aluminum adjuvant (Thermo Scientific, Cat. No. 77161) at a final volume ratio of 1:1 with the immunogen. The mixture was administered via intramuscular injection at a volume of 100 μl to hamsters aged 70-80 weeks, with a booster immunization conducted 21 days later. For the control group (mock), an equivalent volume of PBS mixed with the aluminum adjuvant was processed in the same manner and injected into hamsters of the same age group. Two weeks following the booster immunization, the vaccinated hamsters were challenged intranasally with 10^6^ TCID_50_ of RSV. Post-challenge, the animals were observed daily for clinical signs and weighed. Blood samples were collected via orbital bleeding on days 1, 3, 5, and 7 dpi for CBC analysis. On days 3, 5, 7, and 14, the hamsters were euthanized, and samples of blood, lungs, trachea, and nasal turbinates were collected for subsequent analyses.

### RT-qPCR assay for viral RNA analysis

Total RNA was isolated from samples of hamster blood, lung, trachea, and nasal tissues employing the TRIzol reagent (Invitrogen, Carlsbad, CA, USA). Complementary DNA (cDNA) synthesis was facilitated using the RevCertAid First Strand cDNA Synthesis Kit (Thermo Fisher Scientific, USA). For the detection of viral RNA, the RT-PCR assay was conducted utilizing either the QuantiTect Probe RT-PCR Kit (Qiagen, Germany) in strict adherence to the instructions provided by the manufacturers. Quantification was achieved through the construction of a standard curve, derived from a series of ten-fold serial dilutions of a recombinant plasmid containing a known concentration of the target sequence. The sequences of the primers used were as follows: for RSV forward primer, RSV-F (5’-GGCAAATATGGAAACATACGTGAA-3’), for RSV reverse primer, RSV-R (5’-TCTTTTTCTAGGACATTGTAYTGAACAG-3’), and for the RSV-specific TaqMan probe, RSV-TaqProbe (FAM-CTGTGTATGTGGAGCCTTCGTGAAGCT-MGBNFQ) [26].

### Neutralization assay

Sera samples from vaccinated hamsters and mock hamsters were collected throughout the whole period of vaccination and infection, actually on day 0, 21, 35, 42 and 49 post-vaccination. The CPE method was applied to determine the neutralization titer (NT_50_) of hamster sera in infected Hep-2 cells. Briefly, 5.0 × 10^5^/ml Hep-2 cell suspension was added to 96-well plates and incubated overnight at 37 °C in a carbon dioxide incubator containing 5% CO_2_. Thereafter, series of two-fold dilutions of each hamster serum were prepared using DMEM with 2% FBS, and each dilution was mixed with 100 TCID_50_ of RSV virus at a 1:1 volume ratio, followed by incubation at 37 °C for 1 h. Finally, the mixture of sera and virus was placed to prepared Hep-2 cells (8 wells/dilution), and CPE was observed after four consecutive days of culturing. NT_50_ of sera were determined by calculating the highest dilution of serum that prevented infection in 50% of replicate inoculations.

### Histopathological assessment

The hamsters underwent euthanasia, subsequent to which lung and tracheal tissues were harvested. Prior to any further processing, these tissues were preserved in 40 ml of a 10% (v/v) neutral buffered formalin solution for a duration of seven days to ensure adequate fixation. Following fixation, the tissues were then embedded in paraffin wax, and sections were prepared for staining. These sections were treated with hematoxylin and eosin stains to facilitate histological examination. Visualization of the tissue sections was accomplished using the PANORAMIC 1000 system (3DHISTECH), with subsequent analysis conducted via the CaseViewer 2.4 software (3DHISTECH). The pathological scoring of lung tissue samples is based on six key histopathological features: peribronchial and peribronchiolar inflammatory cell infiltration; bronchiolar epithelial cell injury and desquamation; bronchiolar mucus plugging; perivascular inflammatory cell infiltration; alveolar septal widening and inflammatory cell infiltration; interstitial fibrosis and consolidation. A four-tier grading system is utilized to assess the severity of the histopathological changes: **Grade 0,** Within normal limits. The tissue is considered normal given the animal’s age, sex, and strain under the study conditions. **Grade 1**: Very mild. Changes are just beyond the normal range. **Grade 2**: Mild. Lesions are observable but not severe. **Grade 3**: Moderate. Lesions are prominent and likely to be more severe. **Grade 4**: Severe. Lesions are extremely severe, occupying the entire tissue or organ. The cumulative pathological score for each lung tissue sample was obtained by summing the grades for all six features, resulting in a total score ranging from 0 to 24.

### Immunofluorescence staining

Lung tissue sections embedded in paraffin were subjected to deparaffinization through three successive xylene washes, each lasting 10 minutes. This was followed by a graded hydration process utilizing ethanol concentrations of 100%, 95%, 80%, and 70%. Antigen retrieval was facilitated by heating the sections in 0.01 M citrate buffer, pH 6.0. Subsequent to antigen retrieval, the sections were then blocked with 10% donkey serum or goat serum for 1 hour to prevent nonspecific binding. Overnight incubation at 4°C was performed with primary antibodies, specifically goat anti-RSV (Sigma-Aldrich, ab1128), rabbit anti-EPX (Bioss, bs-3881R) or rabbit anti-MPO (Abcam, ab208670). After incubation with the primary antibody, the sections were treated with Alexa Fluor® 488-conjugated donkey anti-goat IgG secondary antibodies (Abcam, ab150129) or FITC-conjugated goat anti-rabbit IgG secondary antibodies (Abcam, ab6717) for specific detection. The stained coverslips were mounted onto glass slides using a mounting medium containing DAPI to counterstain the nuclei. Fluorescence images were acquired using a confocal microscope.

### RT-qPCR assay for cytokine RNA analysis

cDNA synthesis and quantitative PCR were performed using the One Step TB Gre en® PrimeScript™ RT-PCR Kit (Takara Bio Inc., Shiga, Japan). RT-qPCR reactions were set up in a total volume of 20 μL containing PrimeScript RT Enzyme Mix, TB Green Premix Ex Taq II, RNA template, and gene-specific primers. Thermal cycling conditions consisted of initial reverse transcription at 42°C for 5 min, followed by enzyme activation at 95°C for 10 s, and 40 cycles of denaturation at 95°C for 5 s, annealing at 60°C for 34 s. Fluorescence signals were monitored using a real-time PCR instrument (Applied Biosystems 7500). Relative gene expression levels we re calculated using the comparative Ct method (-ΔCt) with normalization to an endogenous reference gene (γ-actin). The sequence used are as follows: IL-2: forward, 5’-ATGTACAGCAKGCAGCTCGC-3’; reverse, 5’-TGTTGAGATGRYRCTTTGAC-3’; IL-4: forward, 5’-ACAGAAAAAGGGACACCATGCA-3’; reverse, 5’-GAAGCCCTGCAGA TGAGGTCT-3’; IL-5: forward, 5’-GCCGTAGCCATGGAGATC-3’; reverse, 5’-CGATG CACAGCTGGTGGTGAT-3’; IL-10: forward, 5’-GGTTGCCAAACCTTATCAGAAATG-3’; reverse, 5’-TTCACCTGTTCCACAGCCTTG-3’; IL-13: forward, 5’-AAATGGCGGGT TCTGTGC-3’; reverse, 5’-AATATCCTCTGGGTCTTGTAGATGG-3’; IFN-γ: forward, 5’ - TGTTGCTCTGCCTCACTCAGG-3’; reverse, 5’-AAGACGAGGTCCCCTCCATTC-3’; γ-actin: forward, 5’-ACAGAGAGAAGATGACGCAGATAATG-3’; reverse, 5’-GCCTGA ATGGCCACGTACA-3’ [27, 28].

### Quantification and statistical analysis

Data analyses and statistical evaluations were conducted utilizing GraphPad Prism software, version 9. Detailed statistical information pertaining to each experiment is delineated within the respective figure legends. Data representation on both linear and logarithmic scales is articulated as the mean ± standard error of the mean (SEM). Determination of statistically significant disparities was achieved through the application of unpaired t-tests or two-way analysis of variance (ANOVA). A threshold for statistical significance was established at P values less than 0.05. Notations for significance levels are as follows: *P<0.05, **P<0.01, ***P<0.001, and ****P<0.0001, respectively.

## Results

### 1. Clinical symptoms of hamsters following RSV infection

To explore the susceptibility of hamsters to RSV, a range of hamsters of varying ages were infected with RSV. Surprisingly, adult and old hamsters infected with RSV exhibited symptoms including coughing and wheezing between 4 to 8 days post-infection (dpi), which closely resembled the clinical symptoms observed in humans with RSV (Fig. 1A). There was no significant weight loss observed in hamsters across all age groups following infection with either subtype of RSV (Fig. 1B-D). However, we noted that neonatal hamster infected with RSV experienced a markedly slower rate of weight gain compared to the negative control group (Fig. 1B), whereas adult hamsters showed a slight reduction in weight gain rate, albeit not significantly different from the control group (Fig. 1C). The trend of weight change of old hamsters after infection was found to align closely with that of the control group (Fig. 1D). This suggested that RSV infection merely affected weight gain in hamsters, except for the neonatal hamsters during their growth phase.

**Fig. 1.**
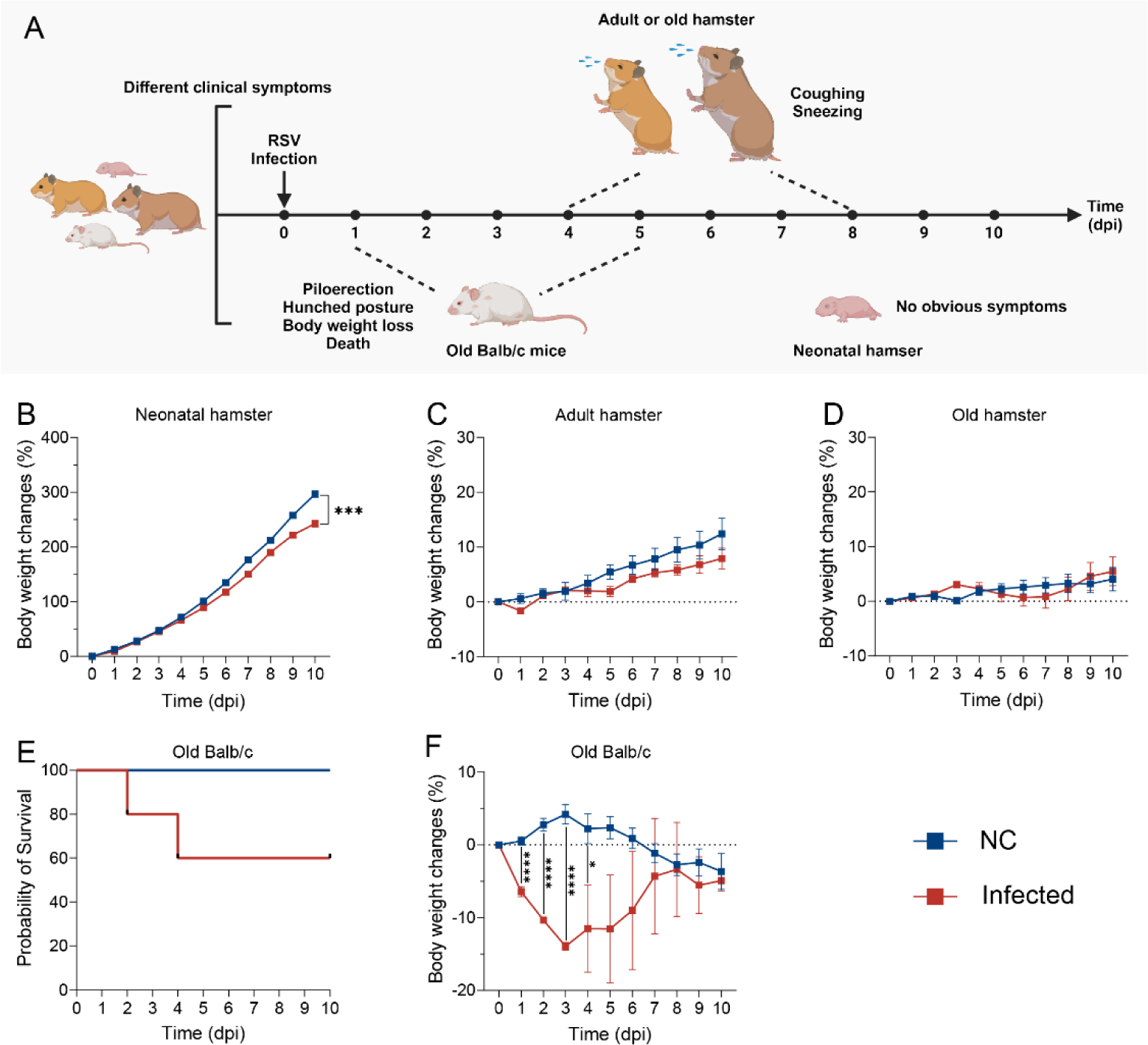
Clinical symptoms and weight changes in Hamsters of different ages following infection with RSV. (A) Illustration of the clinical symptoms and weight changes in hamsters of different ages after RSV infection. (B-D) Weight changes in different age groups of hamsters following RSV infection. (***P < 0.001). (E) Survival probability of old Balb/c mice challenged with RSV. (F) Weight changes in old Balb/c mice following RSV infection (*P<0.05 and ****P<0.0001). Each group consisted of 5 animals.

In the case of old Balb/c mice infected with RSV, clinical signs such as hunched posture, piloerection, and weight loss were observed from 1 dpi to 5 dpi (Fig. 1A). Two mice in the infected cohort died on the 2 dpi and 4 dpi, respectively (Fig. 1E). All mice experienced a notable loss of weight in the initial three days post-infection, with a partial weight recovery observed from 4 dpi (Fig. 1F). By the 7 dpi, the surviving mice had regained weight to levels comparable with those of the negative control group.

Overall, while RSV infection impacted the rate of weight gain in neonatal hamsters, coughing and wheezing were more apparent clinical signs in adult hamsters. Old hamsters infected with RSV showed clinical manifestations highly similar to those observed in humans with RSV, whereas Balb/c mice infected with RSV exhibit typical symptoms of viral infections seen in rodent models.

### 2. Vira replication detected across respiratory tract in hamsters post RSV infection

To ascertain the extent of RSV viral replication in the respiratory tract, lung, trachea, and nasal turbinate tissues were collected at 3 dpi, 5 dpi and 7 dpi in order to determine the viral loads. The results confirmed the successful replication of RSV within the lung, trachea, and nasal turbinate tissues, with viral loads peaking at 3 and 5 dpi (Fig. 2A). Specifically, viral RNA loads in lung tissues reached concentrations of 10^4^ to 10^5^ copies/mg, while nasal turbinate tissues exhibited even higher levels, ranging from 10^5^ to 10^6^ copies/mg. Viral loads within the tracheal tissues were observed to be lower than those observed in the lung or nasal turbinate, with concentrations approximately spanning from 10^3^ to 10^4^ copies/mg. A notable reduction in viral loads across all tissues was observed by 7 dpi (Fig. 2B-D). Given their smaller size, it’s improper to compare neonatal hamsters directly with adult and old ones with regard to the inoculation dose. However, it’s noteworthy that the virus exhibited a greater replication propensity in the upper respiratory tract of neonatal hamsters, as evidenced by higher viral loads accompanied by a distinct delay in virus elimination in the nasal turbinate, rather than in the lung (Fig. 2A). Correspondingly, the lower respiratory tract, especially the lung, was the primary site for viral replication in adult and old hamsters, represented by relatively higher and more resistant viral loads (Fig. S1).

**Fig. 2.**
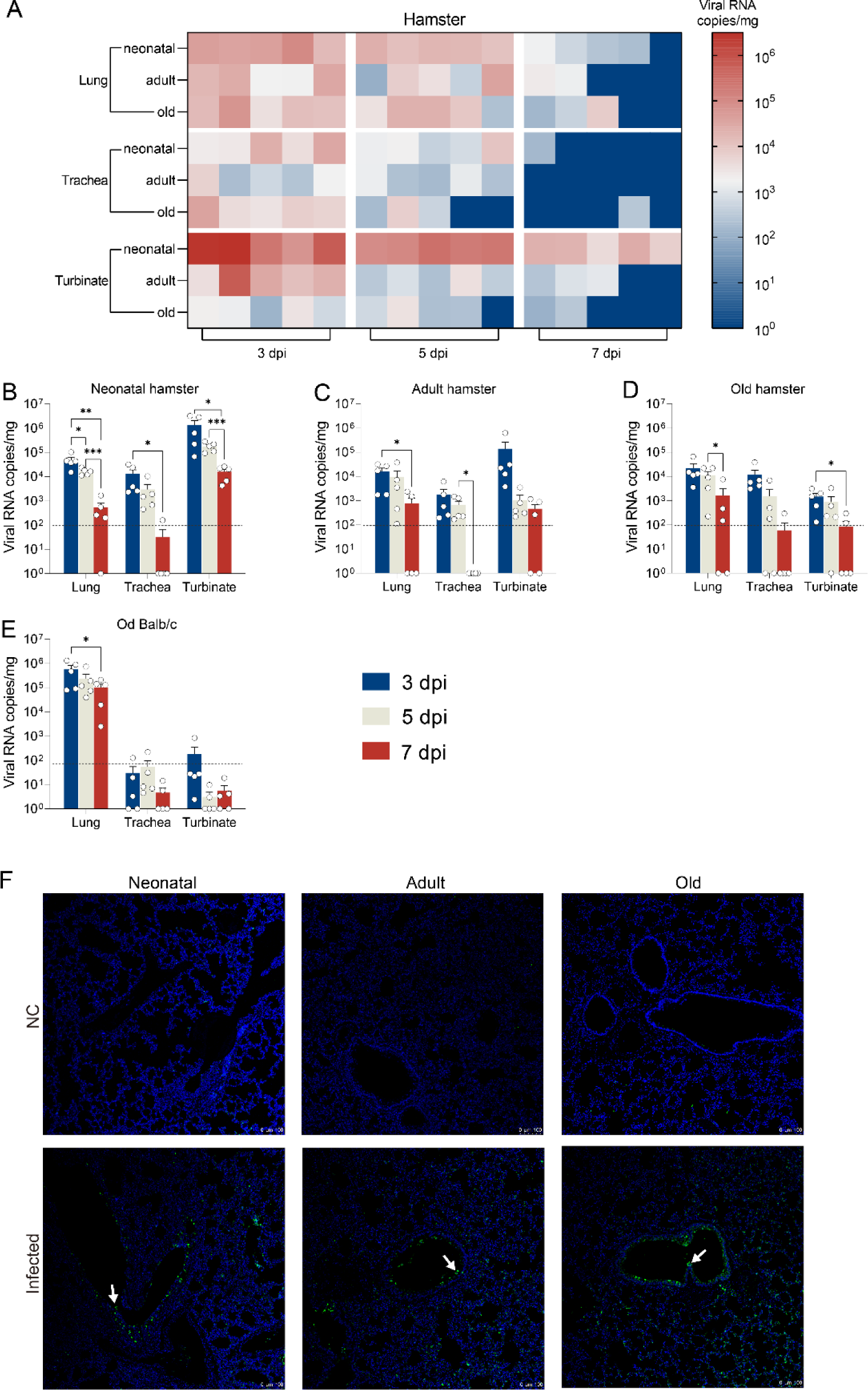
The distribution of the virus in tissues after RSV infection in hamsters. (A) Heat map showing the distribution of viral RNA copies per milligram of tissue in the lungs, trachea, and turbinates of neonatal, adult, and old hamsters infected with RSV. Higher viral RNA loads are indicated by red (higher values), and lower viral loads by blue (lower values). (B-D) Quantification of viral RNA copies of tissue in RSV infected hamsters. (E) Quantification of viral RNA copies of tissue in RSV infected old Balb/c mice. (F) Distribution of viral antigens in hamster lungs post RSV infection. White arrows indicate areas with viral antigen distribution. Each timepoints consisted of 5 animals.

Upon infection with RSV, the viral loads in different tissues of old Balb/c mice exhibited characteristics that differed from those observed in hamsters. Notably, viral concentrations within the pulmonary tissues of Balb/c mice were substantially higher, with values ranging from 10^6^ to 10^7^ copies/mg (Fig. 2D-E). Conversely, viral loads within the tracheal and nasal turbinate tissues of Balb/c mice were found to be significantly lower compared to the lung, with the majority of samples showing viral copy numbers below the detection threshold.

The distribution of RSV viral antigens in lung tissue was examined by immunofluorescence, which revealed that the viral antigens were primarily present in bronchial epithelial cells and were also visible in the alveoli (Fig. 2F). Furthermore, a weaker fluorescent signal was observed in the bronchial lumen, which suggested the presence of desquamated epithelial cells.

In contrast to mouse models, viral replication in RSV-infected hamsters was more evenly distributed throughout the respiratory tract, rather than concentrated primarily in the lungs. This was particularly evident in neonatal hamsters, where viral replication was higher and more persistent in the nasal turbinate. Moreover, the viral antigens were distributed in the epithelial cells of the bronchioles as well as in the alveolar cells in the hamster model. Conversely, in the mouse model, the antigens were primarily concentrated in alveoli.

### 3. Histopathological damage in hamsters upon RSV infection

In order to identify pathological damage to the respiratory tract resulting from RSV invasion, lung and trachea samples were collected for histological examination. In old hamsters infected with RSV, diffuse infiltration of neutrophils was observed in the lungs at 3 dpi, accompanied by minor alveolar dilation and thickening of the alveolar walls, with mild hemorrhage in localized alveolar and bronchial spaces. By 5 dpi, increased neutrophil infiltration was noted, with lymphocytes encircling multiple blood vessels and small bronchioles. The bronchial epithelial cells appeared signs of fragmentation, shedding, and vacuolization, and eosinophilic mucous was visible in a few bronchial lumens. By 7 dpi, the lung inflammation had notably subsided, with a significant reduction in granulocyte and lymphocyte infiltration. However, some bronchial epithelial hyperplasia and irregular arrangement were still observable (Fig. 3A). The lung pathology observed in adult hamsters was less severe compared to the old group, with a reduction in inflammation and structural damage to the small bronchioles (Fig. 3A-B). The neonatal hamsters infected with RSV showed the mildest inflammation, characterized primarily by neutrophil infiltration, with relatively mild bronchial epithelial lesions (Fig. 3A).

**Fig. 3.**
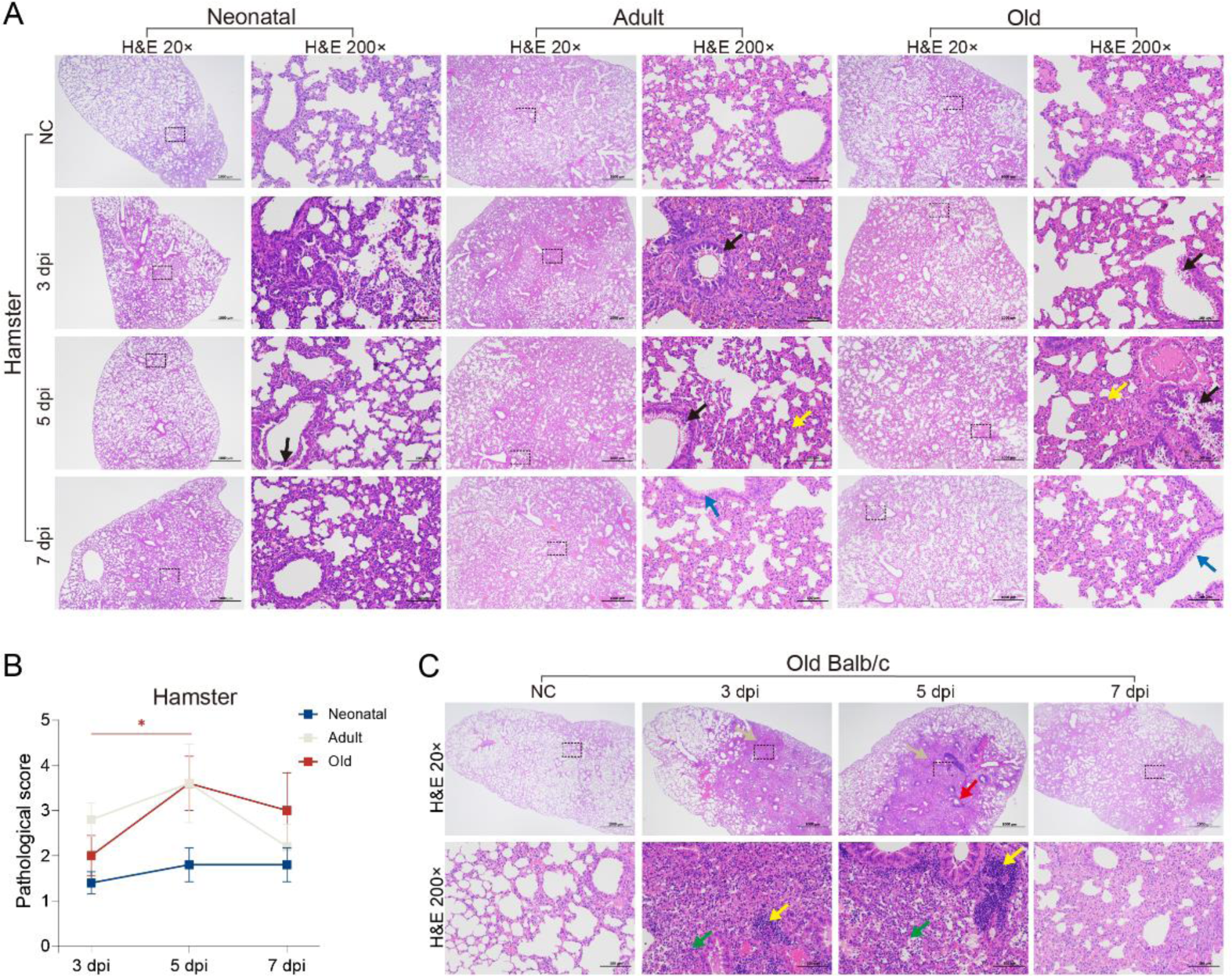
Histopathological analysis of lung tissues from neonatal, adult, and old hamsters infected with RSV at various time points. (A) Representative histopathological images of lung sections from neonatal, adult, and old hamsters infected with RSV. The arrows indicate areas of notable pathological changes: black arrows for bronchiolar epithelial cell damage, yellow arrows for peribronchiolitis, perivasculitis, or alveolitis, and blue arrows for irregular arrangement or fibrosis of the bronchiolar epithelium. (B) Pathological scores of lung tissues from neonatal, adult, and old hamster infected with RSV. Data are presented as mean ± SEM. Statistical significance is indicated by *p < 0.05. (C) Representative histopathological images of lung sections from old Balb/c mice infected with RSV. The arrows highlight areas of pathological changes: grey arrow for lung consolidation, red arrow for peribronchiolitis, green arrow for alveolitis, and yellow arrow for perivasculitis. The dashed boxed area circled in the low-power field indicates the region shown in the high-power field. Each group consisted of 5 animals.

Old Balb/c mice infected with RSV have exhibited severe inflammation as early as 3 dpi, with the condition worsening by 5dpi. This was characterized by infiltration of granulocytes, lymphocytes, and macrophages in the alveoli, and intensified inflammation around bronchioles and blood vessels, including some lymphocytic infiltration into the bronchial mucosal layer. The severe inflammation led to consolidation of lung parenchyma and the loss of alveolar structure (Fig. 3C). Notably, although the inflammation around the alveoli was much more severe in Balb/c mice than in hamsters, their bronchial structures remained relatively intact (data not shown), without the extensive epithelial vacuolation and shedding observed in hamsters (Fig. S2A). Therefore, lung pathology in mice showed predominantly diffuse alveolar damage.

Following the infection of hamsters with RSV, pathological lung damage reached its most pronounced stage at day 5 dpi, manifesting as lesions of bronchial epithelial cells and bronchiolitis, with infiltrates by neutrophils. The older hamsters displayed the greatest degree of lung damage, while the younger hamsters exhibited the least severe pathology. Although the lung pathology observed in mice was more severe, hamsters typically developed bronchiolitis rather than alveolar inflammation in the mice after the RSV infection.

### 4. Viral replication inhibited in hamsters vaccinated with heat-inactivated RSV virus (HI-RSV) after RSV challenge

In order to ascertain the value of the hamster model in evaluating RSV vaccines, particularly with regard to the inactivated vaccines, we administered two intramuscular injections of HI-RSV combined with aluminum adjuvant to 70-80 weeks-old hamsters at a 21-day interval (Fig. 4A). Two weeks subsequent to the booster dose, the animals were infected with RSV. Concurrently, a mock group was given intramuscular injections of aluminum adjuvant mixed with PBS. All vaccinated animals elicited serum neutralizing antibodies against RSV, with the neutralization titer (NT_50_) reaching a level of around 2^6^ after booster immunization. In particular, the highest NT_50_ of vaccinated animals peaked at over 2^8^ after RSV challenge, significantly higher than the mock group (Fig. 4B). The results implied that vaccination with HI-RSV would provide protection against RSV invasion.

**Fig. 4.**
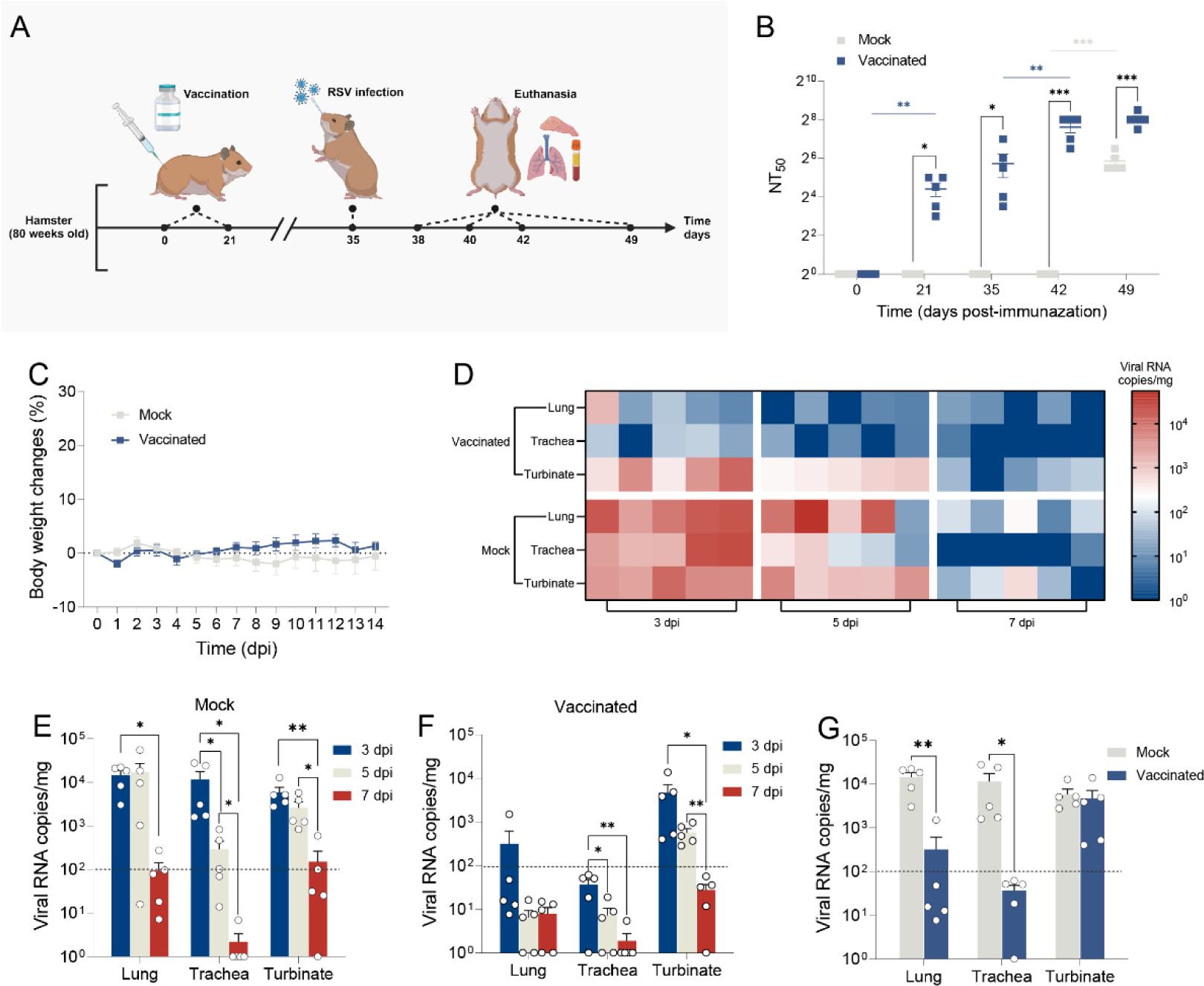
Effects of inactivated vaccination on RSV infection in old hamsters. (A) Schematic representation of the experimental design. 80 weeks old hamsters were vaccinated on day 0, followed by RSV infection on day 35. Euthanasia was performed at different time points post-infection to collect lung tissues for analysis. (B) Neutralizing antibody levels (NT_50_) against RSV in hamster sera from the vaccinated and mock groups at various time points, calculated using the Reed-Muench method. (C) Body weight changes in mock and vaccinated groups over observation period. The percentage change in body weight was monitored daily post-infection. (D) Heatmap showing the viral RNA copies in the lung, trachea, and turbinate tissues of mock and vaccinated hamsters. Color intensity represents the viral RNA copy number, with red indicating higher viral loads and blue indicating lower viral loads. (E) Quantification of viral RNA copies in the lung, trachea, and turbinate tissues of mock hamsters. (F) Quantification of viral RNA copies in the lung, trachea, and turbinate tissues of vaccinated hamsters. (G) Comparison of viral loads in the lungs, trachea, and turbinate tissues between the vaccinated group and the mock group. Data are presented as mean ± SEM. Statistical significance is indicated by *p < 0.05, **p < 0.01 and ***p < 0.001. Each timepoints consisted of 5 animals.

Throughout the 14-day observation period post-infection, no significant differences were observed in the body weight changes between the vaccinated and mock groups (Fig. 4C). However, relatively fewer vaccinated hamsters showed symptoms of coughing (data not shown). Meanwhile, the vaccinated group exhibited markedly lower levels of the viral loads in the lungs and trachea compared to the mock group (Fig. 4D-G). Interestingly, the viral loads in the nasal turbinate showed no significant differences between the vaccinated and mock groups. These findings suggested that the HI-RSV vaccination could inhibit viral replication as well as promote viral clearance from the lungs and trachea in hamsters.

The distribution of viral antigens in the lungs of hamsters also demonstrated the capacity of HI-RSV to impede viral replication. In the vaccinated group, only a minimal quantity of antigen fluorescence was observable, in contrast to the typical viral antigens observed in bronchial epithelial cells in the mock group (Fig. 5B).

**Fig. 5.**
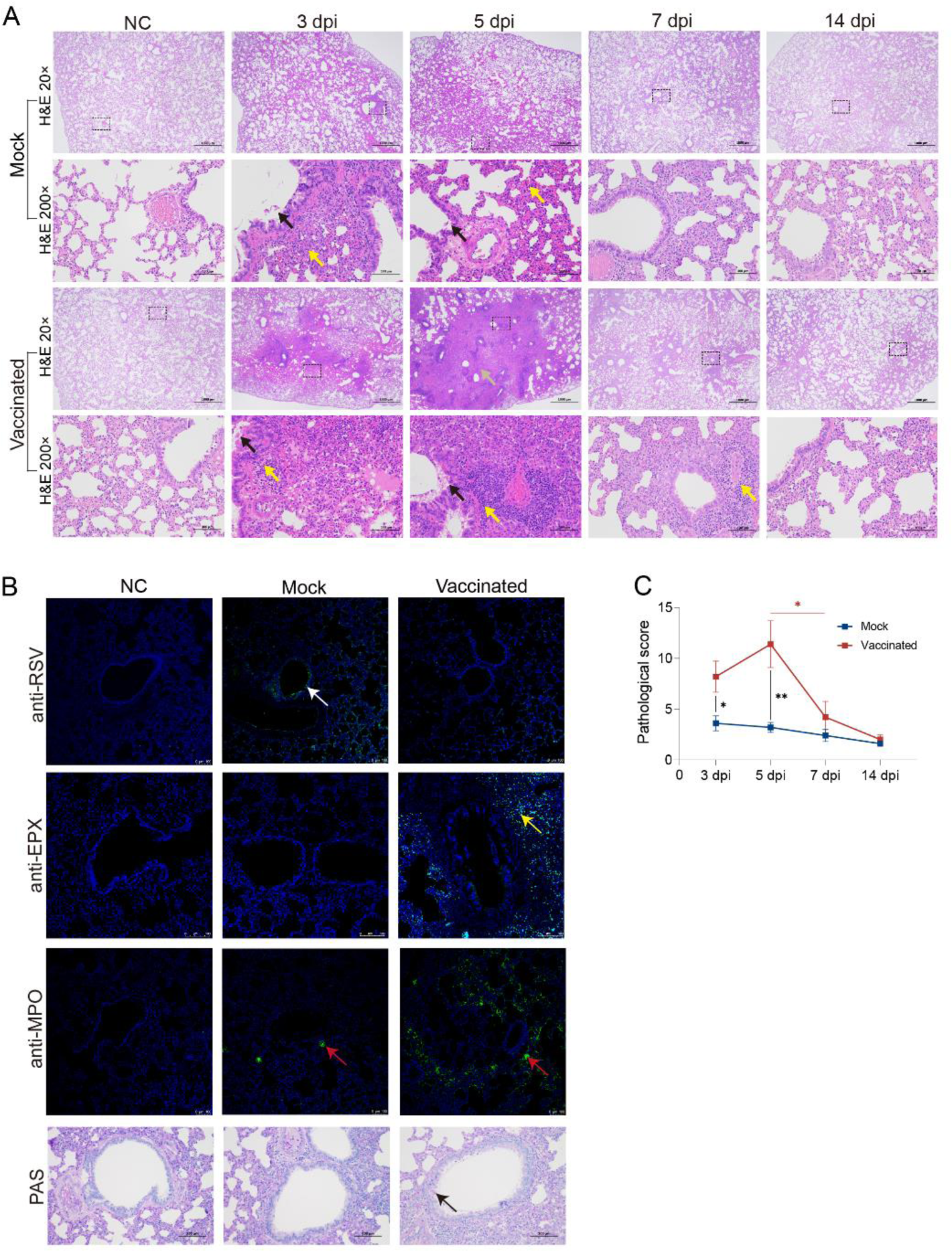
Histopathological and immunofluorescence analysis of lung tissues from mock and vaccinated hamsters infected with RSV. (A) Representative histopathological images of lung sections from mock and vaccinated hamsters. Key pathological features are indicated: black arrow for bronchiolitis, yellow arrows for peribronchiolitis, perivasculitis, or alveolitis and grey arrow for lung consolidation. (B) Immunofluorescence staining of lung sections for RSV antigen, eosinophil peroxidase (EPX) and myeloperoxidase (MPO) in negative control (NC), mock, and vaccinated groups. Arrows indicate areas with positive staining: white arrows for RSV antigen, yellow arrows for EPX and red arrows for MPO. Additionally, Periodic Acid-Schiff (PAS) staining was used to detect mucus production in the lungs, with black arrows indicating mucus-positive areas. (C) Pathological scores of lung tissues from mock and vaccinated hamsters. Data are presented as mean ± SEM. Statistical significance is indicated by *p < 0.05 and **p < 0.01. Each timepoints consisted of 5 animals.

### 5. Severe ERD response induced by vaccination with HI-RSV in old hamsters

Despite the capacity of HI-RSV to neutralize the virus, vaccinated hamsters exhibited more severe pathological lung damage than the mock group (Fig. 5A and C). In fact, the degree of lung damage in the vaccinated hamsters was obviously pronounced at 3 dpi, and continued to worsen, peaking at 5 dpi. It was noted that there had been an increase in the area of tissue consolidation, with unclear alveolar structures and further infiltration by inflammatory cells, including granulocytes and macrophages. The condition of circular lymphocytic infiltration around blood vessels and bronchioles had intensified; more bronchioles contained eosinophilic materials and shed cells, and the epithelial cells exhibited irregular arrangement (Fig. 5A). At 7 dpi, although the lesions had lessened, the alveolar walls still showed substantial granulocyte infiltration, and focal lymphocytic infiltrates persisted around blood vessels and bronchioles (Fig. 5A). By 14 dpi, the lesions had largely resolved, approaching the pathological state of the non-challenged control group (Fig. 5A). As for tracheal lesions, there was no difference observed between vaccinated and mock animals (Fig. S2B).

Subsequently, we employed the use of two antibodies, an anti-eosinophil peroxidase antibody (anti-EPX) and an anti-myeloperoxidase antibody (anti-MPO), to assess the infiltration of eosinophils and neutrophils. Compared to the mock group, a substantial infiltration of eosinophils as well as neutrophils was noticed in the vaccinated group, particularly in the vicinity of bronchioles with an inflammatory response and in areas of consolidation, presenting the strongest fluorescent signals (Fig. 5B). Furthermore, we also performed a periodic acid-Schiff (PAS) staining to detect polysaccharide synthesis in bronchial epithelial cells, and identified an enhanced mucous secretion in the bronchioles of the vaccinated group (Fig. 5B).

Overall, the vaccination with HI-RSV triggered severe ERD, characterized by enhanced pulmonary pathology, extensive eosinophil and neutrophil infiltration and increased mucous secretion in the bronchioles. The extensive damage caused by the HI-RSV overpowered its inhibitory and neutralizing effects against RSV infection, which closely mirrored the pathological conditions observed in clinical cases of humans.

### 6. Immunological response characterized in vaccinated hamsters with ERD

Pulmonary eosinophilia is regarded as a marker of ERD, concurrently, we conducted a complete blood count (CBC) to determine the change of eosinophils in the peripheral blood (Fig. 6). As anticipated, the number as well as the proportion of eosinophils in the vaccinated group were higher than the mock group, particularly at 5 dpi (Fig. 6B and 6E). It is noteworthy that neutrophils in the blood of the vaccinated group demonstrated a tendency to decrease in correlation with the increased neutrophil infiltration observed in the lung tissue, with a significant decline in count occurring at 3 dpi and a significant decrease in the ratio occurring at 5 and 7 dpi. Meanwhile a notable elevation in the number of monocytes was in the vaccinated group was noticed in comparison to the mock group at 5 dpi. Nevertheless, the increase in ratio was observed at as early as 3 dpi, although not reaching a statistically significant difference (Fig. 6C and 6F). This corroborated our pathological findings of an acute neutrophilic infiltration in the lungs post-RSV infection. In addition, a significant decrease in both white blood cell and lymphocyte counts was observed in the vaccinated group at 3 dpi, while the mock group also experienced a slight decline (Fig. S3). These dynamics may be the consequent of the migration of immune cells from peripheral blood to the lung, where viral clearance was initiated.

**Fig. 6.**
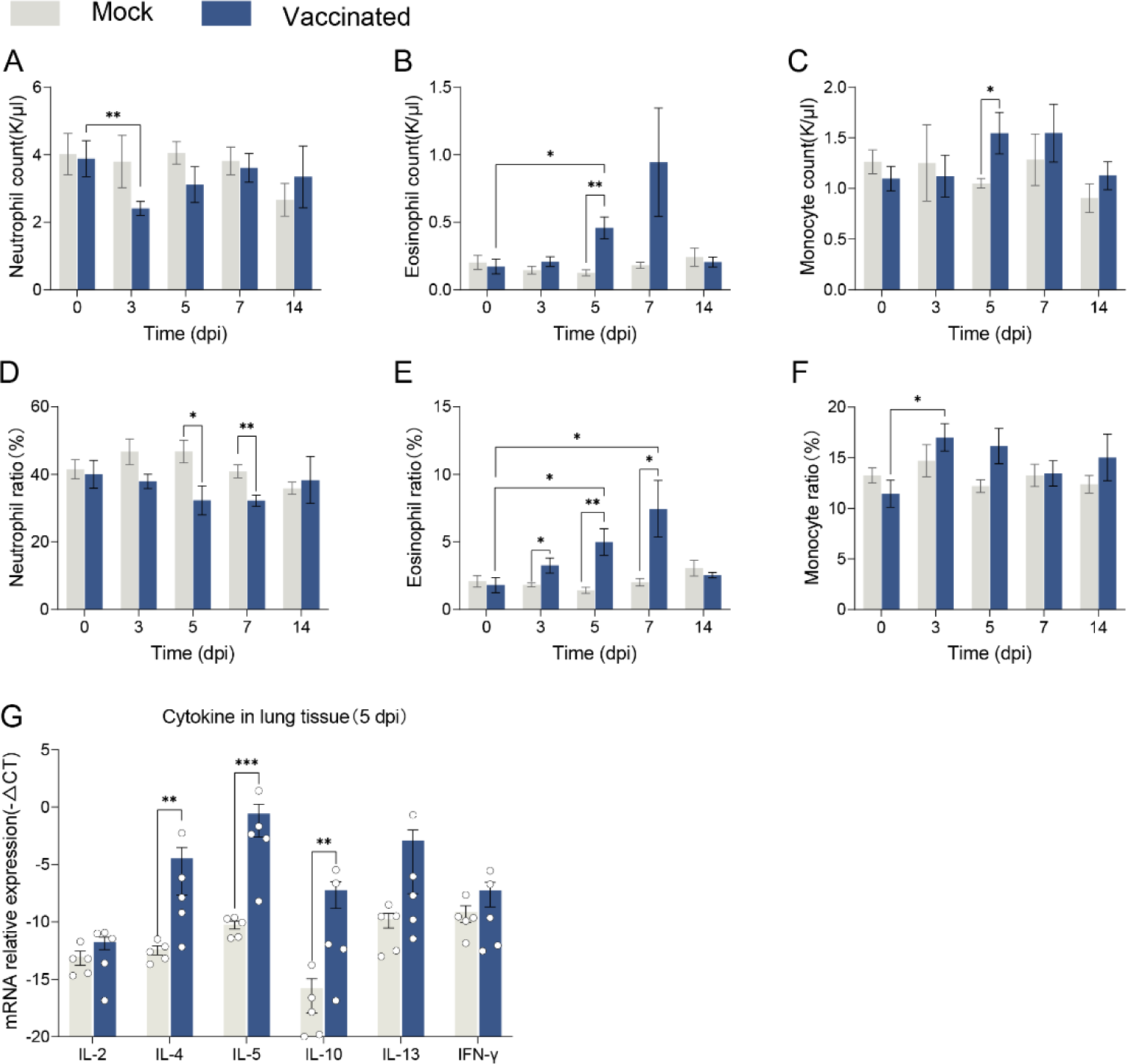
Comparative analysis of blood cell and lung cytokine levels in mock and vaccinated hamsters post-RSV infection. (A-F) Various blood cell parameters were measured in mock (grey) and vaccinated (blue) hamsters. Data are presented as mean ± SEM. (A) Neutrophil count (K/μL). (B) Eosinophil count (K/μL). (C) Monocyte count (K/μL). (D) Neutrophil ratio (%). (E) Eosinophil ratio (%). (F) Monocyte ratio (%). (G) Relative mRNA expression levels of various cytokines in the lungs at 5 dpi. Relative expression levels are represented as -△Ct, with γ-actin as the reference gene. Statistical significance is indicated by *p < 0.05, **p < 0.01 and ***p < 0.001. Each timepoints consisted of 5 animals.

A shift to Th2 response is thought to be one of the key factors in FI-RSV vaccine-induced ERD. Therefore, the transcript levels of the cytokines IL-4, IL-5, IL-10, and IL-13, characteristic of the Th2 response, and the cytokines IFN-γ and IL-2, characteristic of the Th1 response, were determined in lung tissues. Compared to the mock group, the vaccinated group exhibited significantly elevated mRNA levels of IL-4, IL-5, and IL-10. While the increase of IL-13 failed to be statistically significant. On the contrary, the mRNA levels of IL-2 and IFN-γ were comparable between the two groups (Fig. 6D). These results suggested a potential Th2-polarized immune response in the vaccinated hamsters, similar to the characteristics observed in ERD in humans and Balb/c mice [29–32].

In conclusion, the significant changes of eosinophils and neutrophils in the lung and blood, accompanied by Th2-biased response, are indicative of a typical ERD response in HI-RSV immunized hamsters.

## Discussion

RSV significantly threatens the health of infants, young children, and the elderly, and imposes a substantial burden on global health infrastructures [2, 33]. Recent advances in the development of vaccines and therapeutics against RSV have triggered an increase in preclinical studies, consequently elevating the demand for the preferred rodent models for RSV research - cotton rats and mice [34]. However, the clinical symptomatology of both models of RSV infection differs from those of human disease. Moreover, the distinctions between the innate immune response of mice and humans, as well as T cell-specific responses to non-viral antigens of cotton rats, all have an impact on the applicability of the models [20, 35]. Expanding the availability of sophisticated models of RSV infection is always demanding. Syrian hamsters, with greater susceptibility to numerous human pathogens, have been applied for RSV infection, though the comprehensive research in this area remains deficient (24, 25). In order to address the aforementioned knowledge gap, we have developed a Syrian hamster model that has been tested across different age groups of hamsters.

To our surprise, the results demonstrated that RSV infection in both adult and old Syrian hamsters exhibited similar clinical symptoms to those observed in humans, including coughing and wheezing. Concurrently, active viral replication was observed throughout the respiratory tract in hamsters, irrespective of age. While the degree of viral replication was found to be lower in the hamster lungs compared to the Balb/c mice, higher levels of viral load were detected in both the nasal turbinates and trachea of hamsters. Moreover, the lung pathology observed in these hamsters exhibited characteristics consistent with typical viral bronchiolitis, which represents a definitive indicator of RSV infection. The lung lesions in hamsters displayed a closer correlation with the inflammatory response observed in the lungs of RSV patients, which mainly centered on bronchioles, rather than the diffuse alveolar inflammation typically observed after infection in mice [36]. These findings suggested that the hamster model of RSV infection could be a valuable tool for studying RSV. It is capable of replicating both the clinical symptoms and pathological manifestations observed in patients, and thus has the potential to become an important addition to the models currently available.

Hamster of varying ages also possesses a set of distinctive characteristics. Among the three age groups, the old hamsters presented the most severe pulmonary pathological manifestations despite not having the highest levels of viral replication. In contrast, the neonatal hamsters exhibited the highest levels of viral replication, yet displayed the mildest pulmonary pathological manifestations, particularly in the absence of the typical clinical symptoms of coughing. It is possible that the observed discrepancies are attributable to the differing doses of infection, as neonatal hamsters were only capable of receiving one-tenth viral dose. It can be reasonably postulated that neonatal hamsters are inherently less susceptible to RSV. Alternatively, it may also be a consequence of differences in growth and developmental status or immune response between neonatal hamsters and human infants.

The ERD triggered by the FI-RSV vaccine has long been a major concern in the development of RSV vaccines. During a clinical trial in the United States in the 1960s, of the 20 vaccinated children, 16 required hospitalization and two tragically died [37]. Pathological analysis of the lungs from the deceased revealed extensive consolidation due to vigorous inflammatory responses, which resulted in the development of alveolitis and bronchiolitis. In contrast, inflammatory response caused by RSV infection was moderate, without evidence of bronchiolar obstruction or alveolar damage [36, 38–40]. The question of whether the substantial infiltration of eosinophils or neutrophils in the ERD response is a more significant marker is a topic of ongoing debate. The evidence that eosinophils were nearly absent in the RSV-infected patients makes them more distinctive. Concurrently, transcriptional profiles in the lungs of ERD patients also confirmed Th2 response bias, the hypothesis of ERD which derived from mice models [36, 41–43]. It is of the utmost importance to consider the previously mentioned points when establishing ERD animal models, as they represent the fundamental elements in determining and studying ERD responses.

Indeed, the paucity of suitable animal models for the investigation of vaccine-induced ERD effects represents a significant challenge. The ethical concerns for chimpanzees, the sensitization response to non-viral antigens and the absence of eosinophil infiltration in other NHP animals, the lack of exacerbated pulmonary lesions and eosinophil accumulation in newborn lambs, have limited the use of these animal models in ERD research and evaluation [11, 44–46]. As well-established animal models for ERD studies, cotton rat and mouse models were employed not only for the evaluation of vaccine safety but also for comprehensive investigation of ERD mechanisms. Nevertheless, similar to NHP animals, both rodents also produced specific T-cell responses to non-viral antigens in the vaccine, which was further enhanced by RSV antigens [44, 47]. Furthermore, the increase of eosinophils is not inevitable in cotton rat and mouse models. It is therefore prudent to exercise caution when interpreting the results of vaccine candidates evaluated by these models.

In our study, in order to circumvent the potential confounding effects of formalin, hamsters were vaccinated with a heat-inactivated RSV A subtype virus. Following the infection, vaccinated hamsters exhibited alveolitis and bronchiolitis to a significantly greater extent than mock animals, a phenomenon also observed in children who developed ERD. The severe inflammatory response led to substantial interstitial consolidation and fibrosis in the lungs. Furthermore, the aggregated infiltration of eosinophils and neutrophils around the bronchioles, which is indicative of a hallmark event of the ERD response, as well as increased mucus secretion, provided evidences that an ERD response was occurring in the vaccinated hamsters. Although the potential of blood eosinophils as a biomarker for ERD remains uncertain, our observations indicate a consistent increase in their levels within the first seven days post-infection, accompanied by a transient decrease in blood lymphocytes and neutrophils. A subsequent examination of lung tissues further revealed the presence of a Th2 response bias in vaccinated hamsters, which was characterized by the obvious up-regulation of Th2 response signature cytokines, including IL-4, IL-5, and IL-10 as well as the stability of Th1 response signature cytokines, including IL-2 and IFN-γ. In conclusion, the available evidence allows us to reasonably assume that the ERD responses induced by the HI-RSV in hamsters were more severe and distinct than those observed in other rodent models, and closely resembled those observed in humans.

In summary, we have established models using hamsters of different ages infected with two subtypes of RSV. Although different age groups exhibited varying characteristics, their susceptibility to RSV infection renders them a valuable and promising model for further investigation into RSV. The induction of ERD further extended the applicability of the hamster model, including the investigation of RSV pathology, evaluation of vaccine efficacy and safety, as well as the mechanisms of ERD. As a commonly used laboratory animal, the Syrian hamster has a number of advantages, such as strong reproductive ability, smaller space requirements for rearing, and cost-effectiveness compared to the Balb/C mouse. Furthermore, the rapid development of hamster-specific experimental reagents also makes hamster applications more convenient.

## Ethics approval

Animal studies were performed in an animal biosafety level 2 (ABSL-2) facility using high-efficiency particulate air (HEPA)-filtered isolators. All animal procedures were reviewed and approved by the Institutional Animal Care and Use Committee of the Institute of Laboratory Animal Science, Peking Union Medical College (ILAS, PUMC) (LJN-23005).

## Acknowledgment

This work was supported by the National Key Research and Development Project of China (Grant No. 2022YFC2303404), and the CAMS Innovation Fund for Medical Sciences (CIFMS) grant (2022-I2M-1-020).

## Disclosure statement

The authors declare no competing interests.

---

**Fig. S1.**
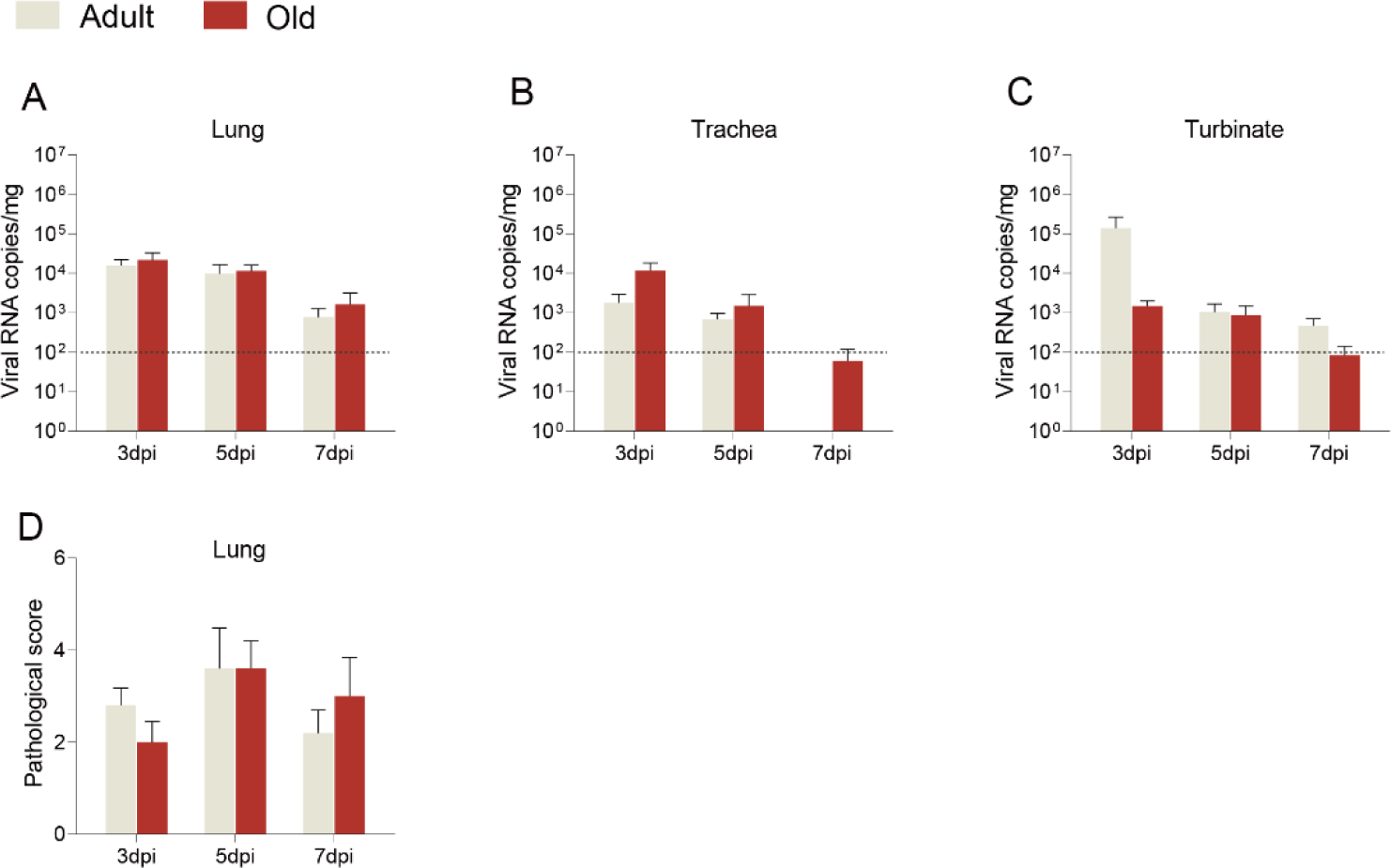
Comparison of viral loads and pulmonary pathological score due to differences in ages. (A-C) Comparison of viral loads in the same tissues of adult (grey) and old (red) hamsters infected with RSV. (D) Comparison of pulmonary pathological score of adult (grey) and old (red) hamsters infected with RSV. Each timepoints consisted of 5 animals.

**Fig. S2.**
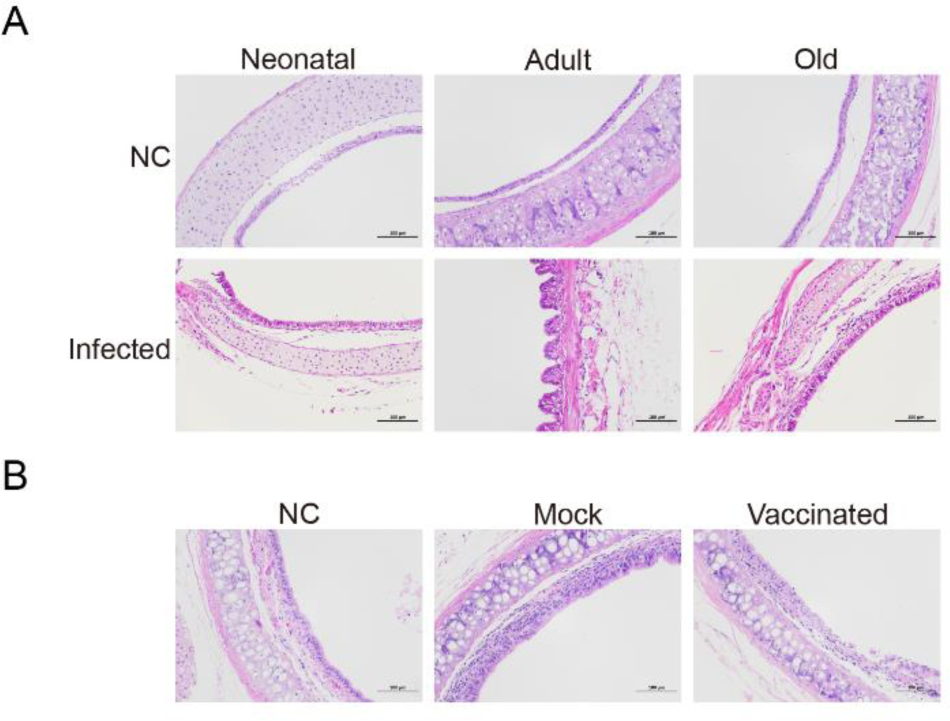
Histopathological analysis of tracheal tissues. (A) Representative histopathological images of tracheal sections from neonatal, adult, and old hamsters infected with RSV at 5 dpi. (B) Representative histopathological images of tracheal sections from negative control (NC), mock, and vaccinated hamsters at 5 dpi. H&E staining was performed and images were captured at 200x magnification. Each timepoints consisted of 5 animals.

**Fig. S3.**
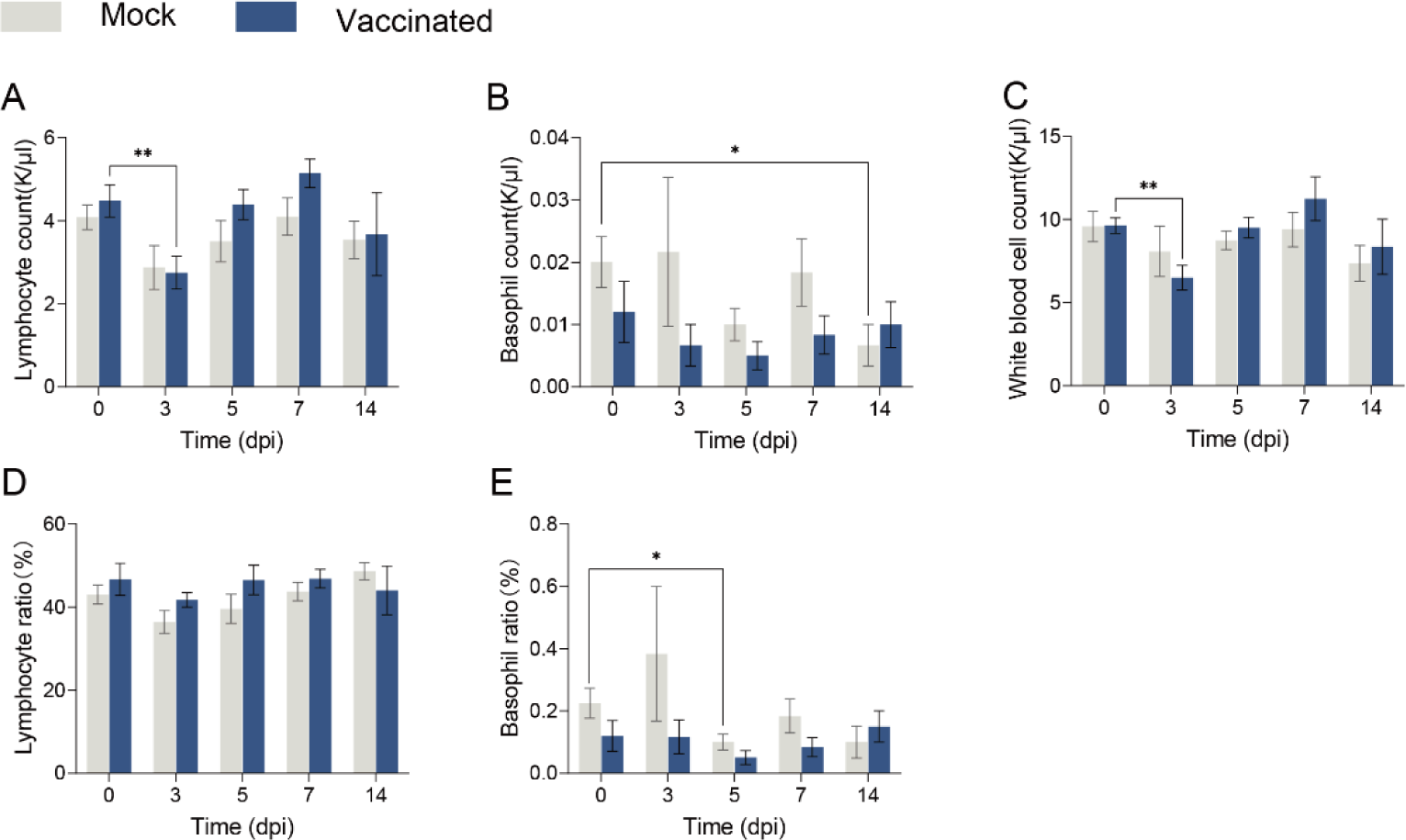
Comparison of changes in white blood cells, lymphocytes, and basophils in the blood of vaccinated and mock groups of hamsters after RSV infection. (A) Lymphocyte count (K/μL). (B) Basophil count (K/μL). (C) White blood cell (WBC) count (K/μL). (D) Lymphocyte ratio (%). (H) Basophil ratio (%).

## References

1. Hacking, D. and J. Hull, Respiratory syncytial virus--viral biology and the host response. J Infect, 2002. 45(1): p. 18–24.

2. Li, Y., et al., Global, regional, and national disease burden estimates of acute lower respiratory infections due to respiratory syncytial virus in children younger than 5 years in 2019: a systematic analysis. Lancet, 2022. 399(10340): p. 2047–2064.

3. Melgar, M., et al., Use of Respiratory Syncytial Virus Vaccines in Older Adults: Recommendations of the Advisory Committee on Immunization Practices - United States, 2023. MMWR Morb Mortal Wkly Rep, 2023. 72(29): p. 793-801.

4. Walsh, E.E., et al., Efficacy and Safety of a Bivalent RSV Prefusion F Vaccine in Older Adults. N Engl J Med, 2023. 388(16): p. 1465–1477.

5. Papi, A., et al., Respiratory Syncytial Virus Prefusion F Protein Vaccine in Older Adults. N Engl J Med, 2023. 388(7): p. 595–608.

6. American Academy of Pediatrics Committee on Infectious, D. and C. American Academy of Pediatrics Bronchiolitis Guidelines, *Updated guidance for palivizumab prophylaxis among infants and young children at increased risk of hospitalization for respiratory syncytial virus infection*. Pediatrics, 2014. 134(2): p. 415-20.

7. Kim, H.W., et al., Respiratory syncytial virus disease in infants despite prior administration of antigenic inactivated vaccine. Am J Epidemiol, 1969. 89(4): p. 422–34.

8. Belshe, R.B., et al., Experimental respiratory syncytial virus infection of four species of primates. J Med Virol, 1977. 1(3): p. 157–62.

9. Taylor, G., Animal models of respiratory syncytial virus infection. Vaccine, 2017. 35(3): p. 469–480.

10. McArthur-Vaughan, K. and L.J. Gershwin, A rhesus monkey model of respiratory syncytial virus infection. J Med Primatol, 2002. 31(2): p. 61–73.

11. Kakuk, T.J., et al., A human respiratory syncytial virus (RSV) primate model of enhanced pulmonary pathology induced with a formalin-inactivated RSV vaccine but not a recombinant FG subunit vaccine. J Infect Dis, 1993. 167(3): p. 553–61.

12. Vaughan, K., G.H. Rhodes, and L.J. Gershwin, DNA immunization against respiratory syncytial virus (RSV) in infant rhesus monkeys. Vaccine, 2005. 23(22): p. 2928–42.

13. Grunwald, T., et al., Novel vaccine regimen elicits strong airway immune responses and control of respiratory syncytial virus in nonhuman primates. J Virol, 2014. 88(8): p. 3997–4007.

14. Wang, D., et al., A Single-Dose Recombinant Parainfluenza Virus 5-Vectored Vaccine Expressing Respiratory Syncytial Virus (RSV) F or G Protein Protected Cotton Rats and African Green Monkeys from RSV Challenge. J Virol, 2017. 91(11).

15. Meyerholz, D.K., et al., Reduced clearance of respiratory syncytial virus infection in a preterm lamb model. Microbes Infect, 2004. 6(14): p. 1312–9.

16. Olivier, A., et al., Human respiratory syncytial virus A2 strain replicates and induces innate immune responses by respiratory epithelia of neonatal lambs. Int J Exp Pathol, 2009. 90(4): p. 431–8.

17. Bem, R.A., J.B. Domachowske, and H.F. Rosenberg, Animal models of human respiratory syncytial virus disease. Am J Physiol Lung Cell Mol Physiol, 2011. 301(2): p. L148–56.

18. Prince, G.A., et al., The pathogenesis of respiratory syncytial virus infection in cotton rats. Am J Pathol, 1978. 93(3): p. 771–91.

19. Taylor, G., et al., Respiratory syncytial virus infection in mice. Infect Immun, 1984. 43(2): p. 649–55.

20. Boukhvalova, M.S., G.A. Prince, and J.C. Blanco, The cotton rat model of respiratory viral infections. Biologicals, 2009. 37(3): p. 152–9.

21. Prince, G.A., et al., Respiratory syncytial virus infection in owl monkeys: viral shedding, immunological response, and associated illness caused by wild-type virus and two temperature-sensitive mutants. Infect Immun, 1979. 26(3): p. 1009–13.

22. Bueno, S.M., et al., Protective T cell immunity against respiratory syncytial virus is efficiently induced by recombinant BCG. Proc Natl Acad Sci U S A, 2008. 105(52): p. 20822–7.

23. Altamirano-Lagos, M.J., et al., Current Animal Models for Understanding the Pathology Caused by the Respiratory Syncytial Virus. Front Microbiol, 2019. 10: p. 873.

24. Rong, N. and J. Liu, Development of animal models for emerging infectious diseases by breaking the barrier of species susceptibility to human pathogens. Emerg Microbes Infect, 2023. 12(1): p. 2178242.

25. Wright, P.F., W.G. Woodend, and R.M. Chanock, Temperature-sensitive mutants of respiratory syncytial virus: in-vivo studies in hamsters. J Infect Dis, 1970. 122(6): p. 501–12.

26. Ho, Y.I.I., A.H. Wong, and R.W.M. Lai, Comparison of the Cepheid Xpert Xpress Flu/RSV Assay to in-house Flu/RSV triplex real-time RT-PCR for rapid molecular detection of Influenza A, Influenza B and Respiratory Syncytial Virus in respiratory specimens. J Med Microbiol, 2018. 67(11): p. 1576–1580.

27. Espitia, C.M., et al., Duplex real-time reverse transcriptase PCR to determine cytokine mRNA expression in a hamster model of New World cutaneous leishmaniasis. BMC Immunol, 2010. 11: p. 31.

28. Ebenig, A., et al., Vaccine-associated enhanced respiratory pathology in COVID-19 hamsters after T(H)2-biased immunization. Cell Rep, 2022. 40(7): p. 111214.

29. Connors, M., et al., Enhanced pulmonary histopathology induced by respiratory syncytial virus (RSV) challenge of formalin-inactivated RSV-immunized BALB/c mice is abrogated by depletion of interleukin-4 (IL-4) and IL-10. J Virol, 1994. 68(8): p. 5321–5.

30. Waris, M.E., et al., Respiratory synctial virus infection in BALB/c mice previously immunized with formalin-inactivated virus induces enhanced pulmonary inflammatory response with a predominant Th2-like cytokine pattern. J Virol, 1996. 70(5): p. 2852–60.

31. Loebbermann, J., et al., Defective immunoregulation in RSV vaccine-augmented viral lung disease restored by selective chemoattraction of regulatory T cells. Proc Natl Acad Sci U S A, 2013. 110(8): p. 2987–92.

32. Knudson, C.J., et al., RSV vaccine-enhanced disease is orchestrated by the combined actions of distinct CD4 T cell subsets. PLoS Pathog, 2015. 11(3): p. e1004757.

33. Li, Z.J., et al., Etiological and epidemiological features of acute respiratory infections in China. Nat Commun, 2021. 12(1): p. 5026.

34. Mazur, N.I., et al., Respiratory syncytial virus prevention within reach: the vaccine and monoclonal antibody landscape. Lancet Infect Dis, 2023. 23(1): p. e2–e21.

35. Shaw, C.A., et al., The role of non-viral antigens in the cotton rat model of respiratory syncytial virus vaccine-enhanced disease. Vaccine, 2013. 31(2): p. 306–12.

36. Polack, F.P., et al., Fatal enhanced respiratory syncytial virus disease in toddlers. Sci Transl Med, 2021. 13(616): p. eabj7843.

37. Kapikian, A.Z., et al., An epidemiologic study of altered clinical reactivity to respiratory syncytial (RS) virus infection in children previously vaccinated with an inactivated RS virus vaccine. Am J Epidemiol, 1969. 89(4): p. 405–21.

38. Russell, C.D., et al., The Human Immune Response to Respiratory Syncytial Virus Infection. Clin Microbiol Rev, 2017. 30(2): p. 481–502.

39. Acosta, P.L., M.T. Caballero, and F.P. Polack, Brief History and Characterization of Enhanced Respiratory Syncytial Virus Disease. Clin Vaccine Immunol, 2015. 23(3): p. 189–95.

40. Joyce E Johnson, R.A.G., Sandy J Olson, Peter F Wright, Barney S Graham, The histopathology of fatal untreated human respiratory syncytial virus infection. Modern Pathology, 2007. 20(1): p. 108–119.

41. Moghaddam, A., et al., A potential molecular mechanism for hypersensitivity caused by formalin-inactivated vaccines. Nat Med, 2006. 12(8): p. 905–7.

42. Boukhvalova, M.S., et al., The TLR4 agonist, monophosphoryl lipid A, attenuates the cytokine storm associated with respiratory syncytial virus vaccine-enhanced disease. Vaccine, 2006. 24(23): p. 5027–35.

43. Sawada, A. and T. Nakayama, Experimental animal model for analyzing immunobiological responses following vaccination with formalin-inactivated respiratory syncytial virus. Microbiol Immunol, 2016. 60(4): p. 234–42.

44. Ponnuraj, E.M., et al., Increased replication of respiratory syncytial virus (RSV) in pulmonary infiltrates is associated with enhanced histopathological disease in bonnet monkeys (Macaca radiata) pre-immunized with a formalin-inactivated RSV vaccine. J Gen Virol, 2001. 82(Pt 11): p. 2663–2674.

45. Clarke, C.J., et al., Respiratory syncytial virus-associated bronchopneumonia in a young chimpanzee. J Comp Pathol, 1994. 110(2): p. 207–12.

46. Derscheid, R.J., et al., Effects of formalin-inactivated respiratory syncytial virus (FI-RSV) in the perinatal lamb model of RSV. PLoS One, 2013. 8(12): p. e81472.

47. Prince, G.A., et al., Vaccine-enhanced respiratory syncytial virus disease in cotton rats following immunization with Lot 100 or a newly prepared reference vaccine. J Gen Virol, 2001. 82(Pt 12): p. 2881–2888.

